# Low-intensity focused ultrasound enables temporal modulation of human midbrain organoid differentiation

**DOI:** 10.1101/2025.08.29.672992

**Authors:** Jinseong Jeong, Yehhyun Jo, Youngsun Lee, Eunyoung Jang, Yeonji Jeong, Won Do Heo, Jennifer H. Shin, Mi-Ok Lee, Hyunjoo J. Lee

**Affiliations:** School of Electrical Engineering, Korea Advanced Institute of Science and Technology (KAIST), Daejeon 34141, Republic of Korea; Stem Cell Convergence Research Center, Korea Research Institute of Bioscience and Biotechnology (KRIBB), Daejeon 34141, Republic of Korea; Department of Bioscience, Korea University of Science and Technology (UST), Daejeon 34113, Republic of Korea; Department of Biological Sciences, Korea Advanced Institute of Science and Technology (KAIST), Daejeon 34141, Republic of Korea; Department of Mechanical Engineering, Korea Advanced Institute of Science and Technology (KAIST), Daejeon 34141, Republic of Korea; Graduate School of Stem Cell and Regenerative Biology, Korea Advanced Institute of Science and y (KAIST), Daejeon 34141, Republic of Korea; KI for Health Science and Technology, Korea Advanced Institute of Science and Technology (KAIST), Daejeon 34141, Republic of Korea; Department of Bio and Brain Engineering, Korea Advanced Institute of Science and Technology (KAIST), Daejeon 34141, Republic of Korea; KAIST Institute for Nano Century (KINC), Daejeon 34141, Republic of Korea

**Keywords:** Focused ultrasound, midbrain organoids, neuromodulation, dopaminergic progenitor cell, differentiation

## Abstract

Controlling the precise timing of biosignaling cues in complex 3D models such as organoids is critical for guiding cellular differentiation and functional maturation. Ultrasound stimulation, a next-generation neuromodulation modality, offers unique advantages due to its target specificity, ability to elicit mechanosensitive responses, and high spatial resolution. However, there is still a lack of precision stimulation platforms and comprehensive biomarker assays for selective control of organoid development. Here, we introduce a modular piezoelectric ultrasound stimulation platform that integrates seamlessly with conventional multi-well plates, enabling selective neuromodulation of midbrain organoids (mBOs). We demonstrate that ultrasound can specifically modulate cellular differentiation by promoting dopaminergic progenitor markers while delaying terminal differentiation. Importantly, ultrasound stimulation did not induce cellular damage, as confirmed by the absence of apoptosis and DNA damage markers. This work demonstrates the potential of focused ultrasound as a safe, non-invasive, and tunable biophysical cue for temporal regulation of organoid differentiation and maturation.

**Significance Statement:** Organoids are three dimensional *in vitro* models derived from human tissue that recapitulate the complex features of organs. However, precise modulation of organoid differentiation relies on biochemical factors, which are limited in their temporal controllability. In this study, we demonstrate that low-intensity ultrasound stimulation enables temporal modulation of midbrain organoid differentiation using a modular multi-well stimulation platform. In particular, we show that ultrasound promotes the proliferation of dopaminergic progenitor cells. These findings suggest that ultrasound can serve as a supplementary mechanical cue to regulate midbrain organoid development.

## Introduction

Human pluripotent stem cell (hPSC)-derived organoids recapitulate the complexity of organs and organ systems, which have enabled critical advancements in developmental studies, drug screening, and direct therapeutic interventions (1–7). The developmental process of these stem cells requires highly specific signaling factors at precise temporal intervals to guide the differentiation and maturation pathways (8–11). Among the early cell types during differentiation, progenitor cells act as a partially specified precursor to a fully differentiated cell type. This intermediate state allows progenitor cells to retain certain cellular characteristics of stem cells with partially determined fates, which is highly beneficial for critical therapeutic applications such as cell transplantation and tissue grafting that require semi-differentiated cells for highly successful integration with the host (12–14). To this end, major advances in biochemical and biophysical signaling techniques, such as bioreactor integration, air-liquid interface cultures, and 3D scaffolding, have contributed to the precise temporal control of progenitor cell differentiation (1, 15–20).

While a biochemical approach forms the basis for organoid development, there is a need for supplementary interventions to overcome the existing challenges of remotely manipulating the differentiation of progenitor cells in functional organoids with high spatiotemporal resolution (8, 10, 21, 22). Direct stimulation modalities such as electrical current, optogenetics, and ultrasound have been used extensively for *in vivo* and *in vitro* neural activation, neuroplasticity modulation, neurogenesis, neuronal differentiation, and cell signaling control (23–33). These techniques have been used in conjunction with conventional biological systems to enable a level of precision not possible with unimodal approaches. Among the direct stimulation modalities, low-intensity focused ultrasound (LIFU) is particularly attractive due to its noninvasiveness, reversible neuronal response, high spatiotemporal resolution, and non-genetic modification (34–46). There have been a couple of studies exploring the effects of ultrasound neuromodulation on cortical organoids; however, these studies have not investigated ultrasound waveform protocols for precise temporal manipulation and are limited in the functional evaluation of various RNA and protein markers of the organoids (47, 48). In addition, there is a lack of high-throughput stimulation arrays for conducting multi-well *in vitro* studies. Moreover, studies on functional organoids that recapitulate distinct brain regions, such as midbrain organoids (mBOs), are necessary due to their association with specific neurological disorders. Investigations into the effect of ultrasound stimulation on cellular functionality, such as neurotransmitter secretion, should be accompanied by the effects on cellular differentiation. Thus, there remains a tremendous challenge to develop and deploy a scalable, multi-well precision ultrasound stimulation platform with comprehensive biomarker analysis for organoids. We hypothesize that critical ultrasound parameters such as pulse repetition frequency (PRF), acoustic intensity, and stimulation duration are able to biophysically modulate the differentiation timing and dopaminergic function of free-floating mBOs.

In this work, we developed a piezoelectric-based ultrasound transducer platform on a modular printed circuit board (PCB), with impedance-matching considerations for robust multi-well stimulation and transducer design considerations for various conventional well plates. Using this system in a 24-well plate, we successfully verified acute and chronic neuronal activation of the mBOs over 55 days without cellular damage. Ultrasound promoted the selective differentiation of dopaminergic progenitor cells without inducing cellular damage. This precise modulation was achieved without altering the overall development of other cell types in the midbrain organoids. Our approach supports the utility of ultrasound as a potential mechanical cue for regulating the temporal differentiation and function of midbrain organoids.

## Results

### Development of the ultrasound platform and stimulation protocol

We designed a modular and robust platform capable of multi-well stimulation for various conventional well plates. We packaged the PUT in a modular platform with a robust encapsulation layer that doubled as an acoustic coupling medium. To accommodate various configurations of well plate shapes and sizes, we designed our system to be modular by fabricating the single-element PUTs on individual printed circuit boards (PCBs) and using a multi-connector board to drive multiple devices simultaneously. This design enabled multi-well stimulation while maintaining the flexibility to configure the devices as single-element units (Figure 1A, Figure S1, Methods).

**Figure 1.**
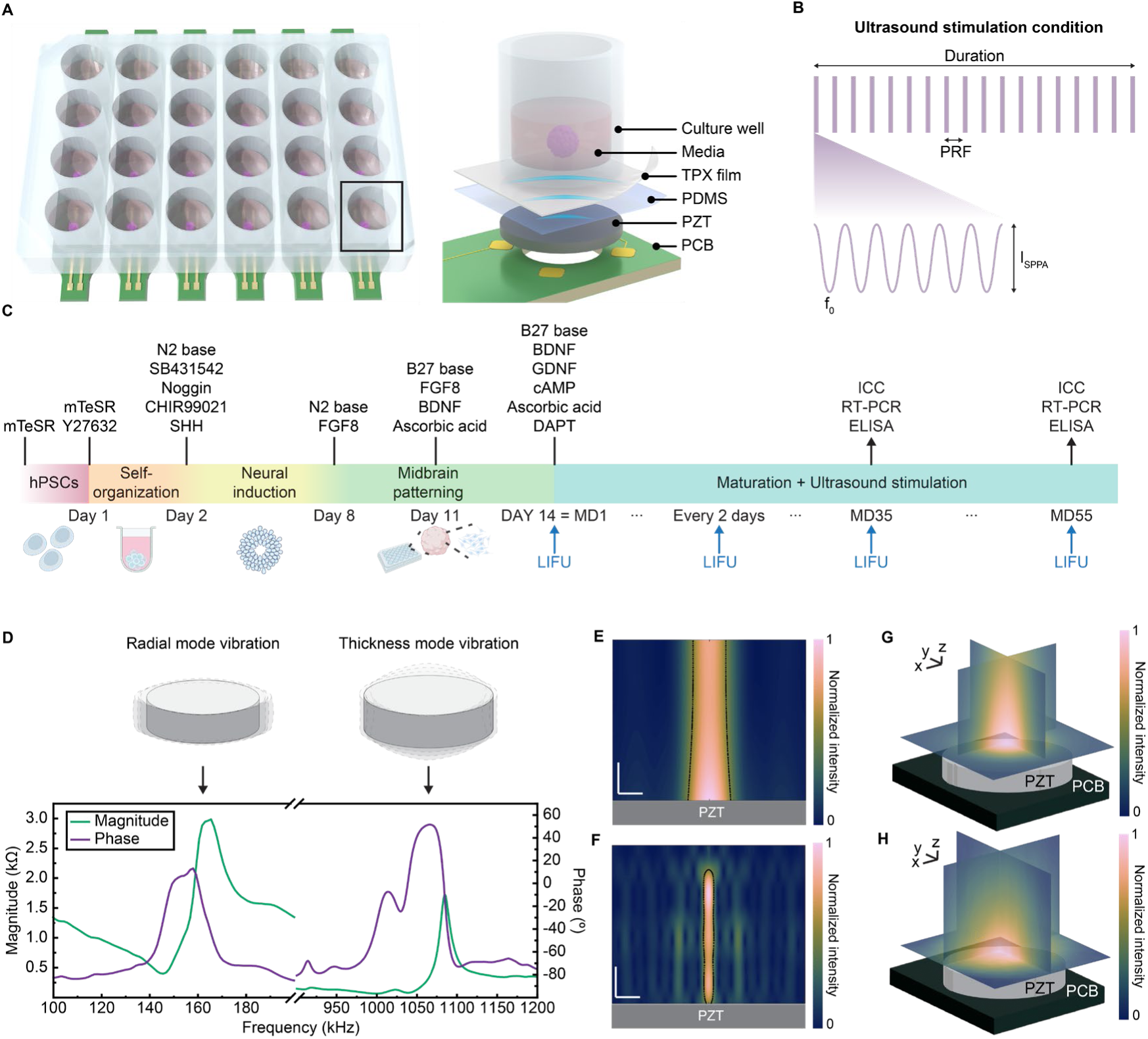
Ultrasound stimulation of midbrain organoids. (A) Schematic illustration of the ultrasound stimulation platform. (B) The ultrasound waveform used to stimulate midbrain organoids. (C) Timeline for generation of midbrain organoids and ultrasound stimulation. (D) Impedance of the two different resonance modes. (E and F) Simulated beam profile generated by operating PZT discs in two different resonance modes. Scale bar, 0.5 mm (vertical) and 2 mm (horizontal). (G) Simulated 3D beam profile produced by operating a PZT disc in the radial resonance mode. Scale bar, 1 mm (x-axis), 1 mm (y-axis), and 0.1 mm (z-axis). (H) Measured 3D beam profile produced by operating a PZT disc in the radial resonance mode. Scale bar, 1 mm (x-axis), 1 mm (y-axis), and 0.1 mm (z-axis).

It is well established that the ultrasound waveform parameters, particularly the PRF, are critical for eliciting cell-type-specific responses (30, 49–53). As an initial parameter screening, we conducted a pilot test for various PRFs. We stimulated midbrain organoids for 7 days with three different PRFs based on existing literature protocol. We evaluated the relative mRNA expression of the neuronal activation marker, *cFOS*, and dopaminergic neuron marker, tyrosine hydroxylase (*TH*), using RT-PCR. Based on these preliminary findings, we selected a PRF for all subsequent experiments.

To demonstrate the temporal effects of ultrasound on the differentiation timing in mBOs, we delivered LIFU stimulation for 10 minutes every two days for 55 days. Based on previously reported results, we hypothesized that LIFU stimulation would effectively stimulate and upregulate neuronal processes of the mBOs (Figure 1B). A control condition with no stimulation was used to measure the effects of ultrasound stimulation.

The midbrain organoids were generated according to the protocol described in the Methods section (Figure 1C, Methods). The hESCs were seeded in ultra-low attachment 96-well plates to facilitate embryoid body formation, and differentiation was initiated by transitioning to N2 medium with SB431542, Noggin, CHIR99021, and SHH. This induction medium was refreshed on days 2, 4, and 6. On day 8, cells were cultured in N2 medium containing FGF8, followed by a switch to B27-based medium on day 11, which was enriched with FGF8, BDNF, and ascorbic acid. From day 14 onward, the organoids were maintained in maturation medium containing BDNF, GDNF, ascorbic acid, db-cAMP, and DAPT, with regular medium changes. Ultrasound stimulation was initiated at maturation day 1 (MD1) and continued every other day until MD55. Organoids were harvested on maturation day 35 (MD35) and maturation day 55 (MD55). ICC, RT-PCR, and ELISA were conducted for MD35 and MD55 organoids (Methods).

For *in vitro* applications using ultrasound stimulation, it is crucial to produce a beam that uniformly encloses the entire biological sample. A full-width half maximum (FWHM) beam size of 2 mm (lateral) by 1 mm (axial) was required to uniformly envelope the mBOs at the bottom of the well plate. To achieve the desired size and location of the ultrasound beam, we considered the resonance modes in the lead zirconate titanate (PZT) disk and the size of the well plates. There are two different resonance modes in the PZT disk: thickness mode and radial mode (Figure 1D). Considering that most PZT disks feature a thin and wide cylindrical geometry (i.e., a small thickness and a large diameter), the radial resonance mode frequency is lower than the thickness resonance mode frequency. These geometries indicate that the operation of the PZT disk in radial mode generates a large lateral FWHM beam right in front of the PZT disk surface due to its low frequency. Using finite element method (FEM) simulations, we confirmed that the radial mode beam profile had a lateral FWHM of 3.6 mm and an axial FWHM of 2.6 mm. In contrast, the thickness mode operation had a lateral FWHM of 0.3 mm and an axial FWHM of 2.1 mm (Figure 1E, F). Therefore, to generate a wide beam, radial mode operation was selected for all subsequent experiments.

To achieve the desired size and location of the ultrasound beam, we simulated various PZT and well dimensions. We also conducted FEM simulations for conventional 6-well, 24-well, and 96-well plates (Figure S3, Methods). We examine two different parameters: the maximum intensity variation within the organoid space and the ratio of the 50% intensity volume within the organoid space to evaluate uniformity. The FEM simulation results of the former parameter were 89%, 75%, and 53% for 6-well, 24-well, and 96-well plates, respectively. The FEM simulation results for the latter parameter were all 100% for the 6-well, 24-well, and 96-well plates. Therefore, it is possible to use any size of well plate for the experiment. In this work, we used a 24-well plate for all experiments because it offers a compact format suitable for high-throughput applications. The resultant beam profile of the 24-well platform operating in the radial mode had a lateral FWHM of 4.9 mm and an axial FWHM of 1 mm. The maximum intensity variation within the organoid space was 72%, and the ratio of the 50% intensity volume within the organoid space was 64%, which was consistent with our FEM simulation results (Figure 1G, H).

We conducted our experiments in a conventional incubator, which was maintained at a high humidity (∼70%) and warm temperature (∼37°C). Thus, robust encapsulation of the stimulation platform with a hydrophobic, elastic, and acoustically transparent material was critical for stable, long-term stimulation. After packaging our PUT on the PCB, we dip-coated the entire device in polydimethylsiloxane (PDMS) and measured the electrical impedance throughout multiple experimental cycles in the incubator (Figure S4, Methods). We confirmed that our device maintained robust operation after nine 6-hour experimental cycles in the incubator. The measured beam profile of six PUTs demonstrated that there was less than 15% variability in the output acoustic intensity (Figure S5, Methods). This level of consistency was sufficient for successful ultrasound stimulation as demonstrated by our *in vitro* results.

### Ultrasound stimulation robustly activates neurons in the organoids

To investigate whether midbrain organoids were stimulated by ultrasound, we evaluated the gene expression levels and protein levels of cFOS and activation of yes-associated protein (YAP). First, we analyzed the change in cFOS, which is an immediate early gene (IEG) widely used as a biomarker for neural activation (54). For the US group, the relative mRNA expression of *cFOS* was significantly increased on both MD35 and MD55 (Figure 2A). In addition to qPCR results, the protein level of cFOS was compared across the experimental groups. The cFOS protein level was significantly increased in the US group on both MD35 and MD55 (Figure 2B-D). These cFOS analysis results implied that ultrasound successfully stimulated the midbrain organoids.

**Figure 2.**
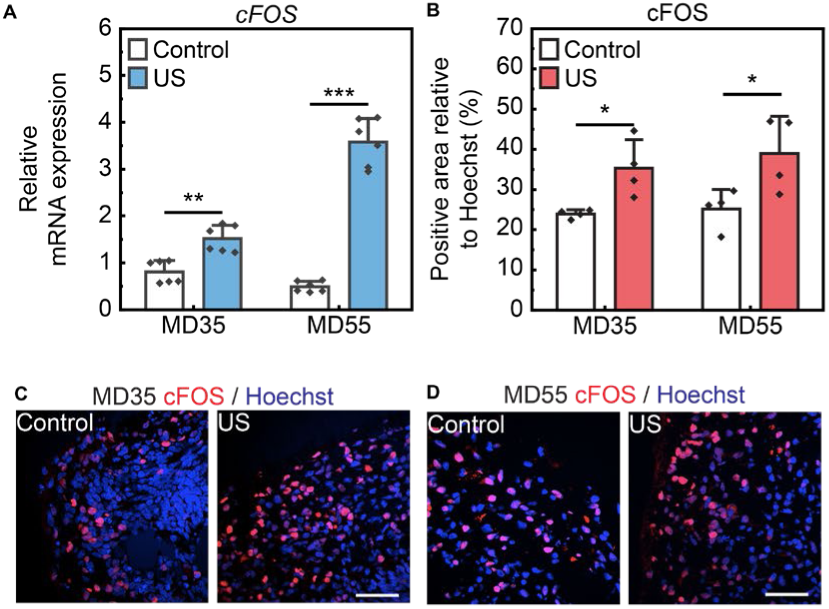
Activation of neural cells by ultrasound stimulation. (A) Relative mRNA expression of *cFOS* in the US group. MD35, Mann-Whitney U test, *U=0, **p<0.01, n=6*. MD55, two sample two-sided unpaired Student’s t-test, *t=-14.70, ***p<0.001, n=6*. (B) Relative protein level of cFOS in the US group. MD35, two sample two-sided unpaired Student’s t-test, *t=-3.210, *p<0.05, n=4*. MD55, two sample two-sided unpaired Student’s t-test, *t=-2.654, *p<0.05, n=4*. (C and D) Representative immunostaining images of cFOS. Scale bar, 50 μm. Solid bars and error bars indicate mean value and standard deviation, respectively.

Next, we observed the translocation of YAP by immunostaining. YAP is one of the transcription coregulators in the Hippo signaling pathway and is activated by various triggers such as the cell density and mechanical cues like the stiffness of the extracellular matrix (ECM). When YAP is activated, phosphorylation of YAP is inhibited, and YAP is translocated into the nucleus. As a mechanical stimulus, ultrasound stimulation is known to be a possible trigger of YAP activation (55, 56). Therefore, we analyzed the YAP ratio in the nucleus and cytoplasm using immunostaining techniques. Representative fluorescence images exhibited higher nuclear translocation of YAP for both MD35 and MD55 in the stimulated group compared to the control (Figure S6A, B). For the US group, we observed a slight increase in the ratio of nuclear to cytoplasmic YAP on MD35, while the MD55 organoids demonstrated significantly more YAP translocated into the nucleus (Figure S6C). We also investigated the relative gene expression of the connective tissue growth factor (*CTGF*), which is the downstream target gene of the YAP/TAZ signaling pathway. When the activated YAP is translocated into the nucleus, YAP/TAZ combines with transcriptional enhanced associate domain (TEAD) transcription factor and transcribes downstream target genes such as *CTGF*, cysteine-rich angiogenic inducer 61 (*CYR61*), and amphiregulin (*AREG*). The expression of *CTGF* was increased in the US group organoids on MD35 (Figure S6D). There was no significant change in *CTGF* in the MD55 organoids. These results confirmed that YAP was activated through both the translocation of YAP and the expression of *CTGF*. Collectively, ultrasound stimulation of midbrain organoids using both conditions successfully activated the mBO cells and regulated the Hippo signaling pathway, which affects the proliferation and differentiation of midbrain organoids. These results verified that ultrasound is capable of neural activation in mBOs, which was in line with the current reports on ultrasound neuromodulation. Based on these results, we proceeded to investigate the functional responses of mBOs to ultrasound stimulation.

### Ultrasound stimulation promotes the induction of dopaminergic progenitor cells

To determine the functional effects of mBO neuromodulation, we investigated the impact of ultrasound stimulation on critical cellular processes such as differentiation and maturation. Ultrasound was delivered from MD1, with the mBOs already guided into the midbrain lineage (Figure S7). Since mBOs primarily consist of dopaminergic cell lines, we first investigated the influence of ultrasound stimulation on dopaminergic progenitor cells. By immunostaining the dopaminergic progenitor marker, Forkhead box protein A2 (FOXA2), representative immunostaining images demonstrated higher levels of FOXA2 in both the MD35 and MD55 organoids in the stimulated group compared to the control (Figure 3A, B). Quantitatively, we verified that FOXA2 levels were significantly increased in organoids stimulated with ultrasound on MD35 and MD55 (Figure 3C). Additional qPCR results demonstrated the relative gene expression level of the LIM homeobox transcription factor 1 alpha (*LMX1A*), which was increased in organoids stimulated with ultrasound on MD35 and MD55 (Figure 3D) These findings suggest that successful ultrasound stimulation causes an increase in the population of dopaminergic progenitor cells. Since ultrasound induced an increase in the population of dopaminergic progenitor cells, we next examined the dopaminergic function of the stimulated mBOs. We analyzed tyrosine hydroxylase (TH), a biomarker for dopaminergic cells, using both protein and gene analysis. First, we evaluated the relative mRNA expression of *TH*. For the US group, we did not observe any significant changes in *TH* expression levels on MD35 (Figure 4A). However, a significant decrease in *TH* gene expression was observed for the US group on MD55. For the TH protein expression levels, there were no significant differences between the control and the US group on MD35 and MD55 (Figure 4B-D).

**Figure 3.**
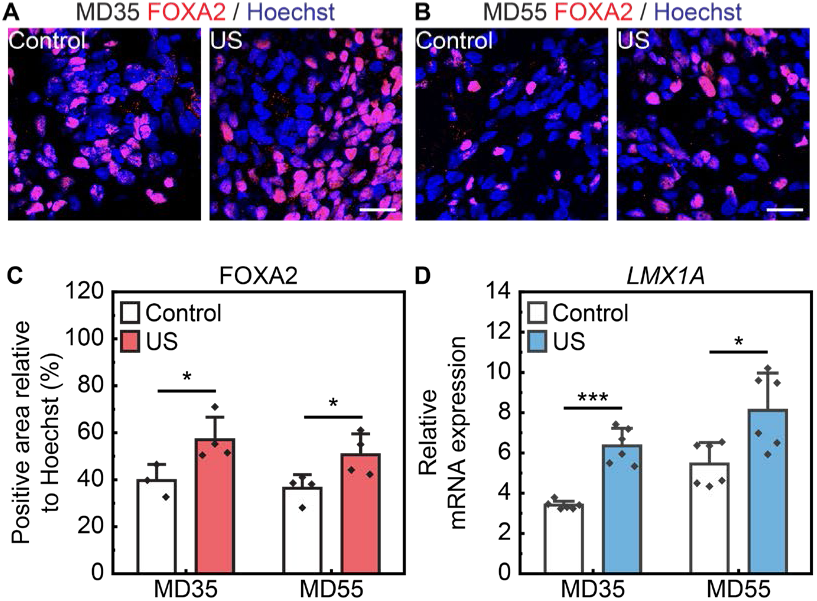
Controlled differentiation of dopaminergic progenitor cells using ultrasound stimulation. (A and B) Representative immunostaining images of FOXA2. Scale bar, 20 μm. (C) Relative protein level of FOXA2 in the US group. MD35, two sample two-sided unpaired Student’s t-test, *t=–2.800, *p<0.05, n=4*. MD55, two sample two-sided unpaired Student’s t-test, *t=–2.674, *p<0.05, n=4*. (D) Relative mRNA expression of *LMX1a* in the US group. MD35, two sample two-sided unpaired Student’s t-test, *t=–7.979, ***p<0.001, n=6*. MD55, Mann-Whitney U test, *U=4, *p<0.05, n=6*. Solid bars and error bars indicate mean value and standard deviation, respectively.

**Figure 4.**
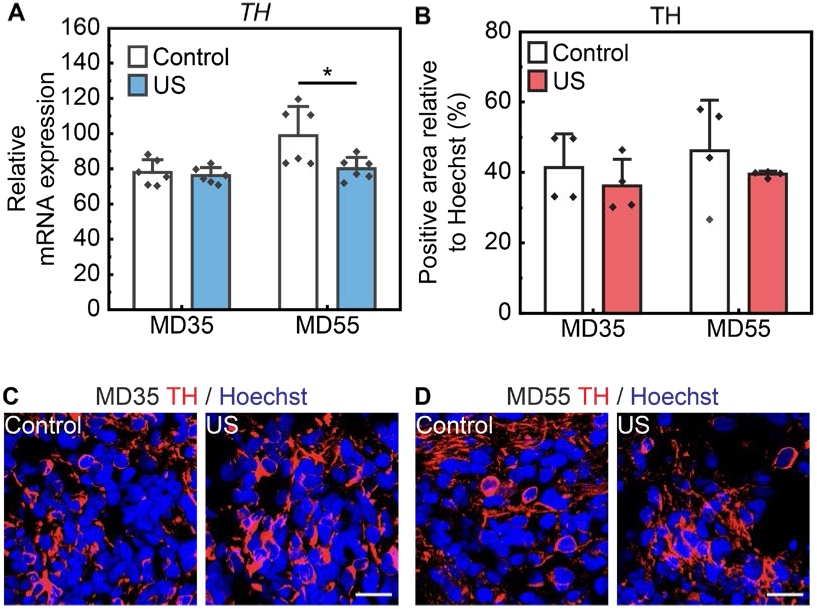
Controlled differentiation of dopaminergic neurons using ultrasound stimulation. (A) Relative mRNA expression of *TH* in the US group. MD35, two sample two-sided unpaired Student’s t-test, *t=0.519, p=0.617, n=6*. MD55, two sample two-sided unpaired Student’s t-test, *t=2.586, *p<0.05, n=6*. (B) Relative protein level of TH in the US group. MD35, Mann-Whitney U test, *U=12, p=0.343, n=4*. MD55, two sample two-sided unpaired Student’s t-test, *t=0.925, p=0.423, n=4*. (C and D) Representative immunostaining images of TH. Scale bar, 20 μm. Solid bars and error bars indicate mean value and standard deviation, respectively.

We also assessed the differentiation of the mBO cells into neurons by evaluating the expression levels of both the microtubule-associated protein 2 (*MAP2*) gene and MAP2, a protein marker for neurons. There was no significant difference in both the mRNA expression level of *MAP2* and the protein expression level of MAP2 for the US group on MD35 and MD55, except for a significant increase in the mRNA *MAP2* expression on MD55 (Figure S8A-D). These findings suggested that the ultrasound-induced increase in dopaminergic progenitor cell population slightly inhibited differentiation into dopaminergic cells. This effect was to be expected considering that the overall size of the stimulated mBOs did not increase compared to the control (Figure S9A, B).

In addition to TH, we quantified the amount of dopamine secretion as a direct barometer of dopaminergic function using an enzyme-linked immunosorbent assay (ELISA). Dopamine levels were slightly decreased on MD35 and MD55 for the US group (Figure S10). These results are consistent with the TH expression data and suggest that ultrasound stimulation under these conditions promotes expansion of the progenitor population while subtly inhibiting dopaminergic differentiation and function. This supports the potential of ultrasound as a noninvasive tool for temporal modulation of early-stage differentiation in organoids.

### Ultrasound stimulation modulates transcriptomic signatures

To confirm the efficient differentiation of hPSCs into mBOs, we conducted transcriptomic comparison between undifferentiated hPSCs and day 35 mBOs. Bulk RNA sequencing data revealed a clear separation between the two groups, with robust induction of neurodevelopmental and region-specific transcriptional programs in mBOs (Figure S11A). Gene ontology (GO) biological process enrichment analysis of differentially expressed genes (DEGs) exhibited significant enrichment in pathways associated with nervous system development, neuron differentiation, regulation of dopamine secretion, and dopamine metabolic process in the mBO groups (control, US) (Figure S11B). KEGG pathway analysis further confirmed the upregulation of dopaminergic synapse, morphine addiction, and oxytocin signaling pathway in mBOs, which are critical for the midbrain (Figure S11C). In contrast, the hPSCs group exhibited high expression of pluripotency and proliferation-associated genes such as MYC and CDC20, and demonstrated enrichment for cell cycle regulation, DNA replication, and stem cell maintenance pathways. These findings validate the successful and developmentally appropriate differentiation of hPSCs into midbrain lineage by day 35. Importantly, the presence of ultrasound stimulation did not disrupt this trajectory, indicating that ultrasound does not compromise differentiation fidelity or lineage identity. To further investigate the transcriptomic impact of ultrasound stimulation on mBOs, bulk RNA sequencing was performed on organoids subjected to ultrasound conditions (US) and compared to non-stimulated controls (control) at MD35, a timepoint corresponding to early-stage lineage modulation. Principal component analysis (PCA) was conducted to assess the global transcriptomic variance across groups (Figure 5A). The 3D PCA plot revealed clear separation among the control and US group samples along the first three principal components (PC1: 42.4%, PC2: 31.9%, PC3: 25.8%). Each group formed a distinct cluster, which indicated that ultrasound stimulation induced condition-specific transcriptomic profiles. Notably, the separation between the control and US groups suggested that ultrasound differentially modulated gene expression programs in mBO. In the US group, 102 genes were upregulated, and 75 genes were downregulated (Figure 5B).

**Figure 5.**
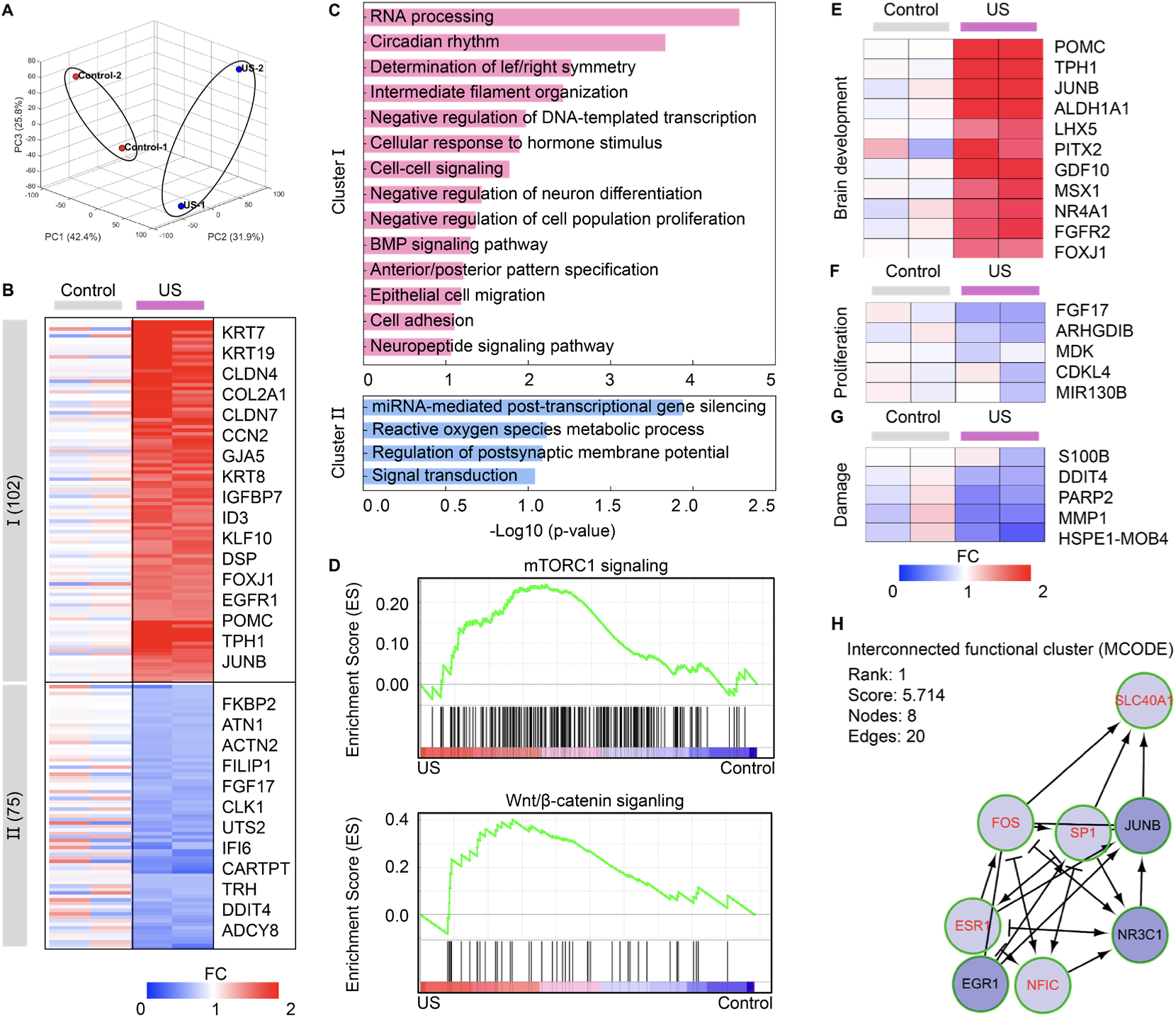
Total transcriptome analysis. (A) 3D PCA plot of two different groups. (B) Heatmap plot of DEGs. (C) GO analysis results of each cluster. (D) Enrichment plot of the mTORC1 signaling and WNT/β-catenin signaling in the US group. (E) Heatmap of genes related to brain development. (F) Heatmap of genes related to the proliferation. (G) Heatmap of genes related to the damage. (H) MCODE analysis of the US group.

Gene ontology (GO) enrichment analysis of differentially expressed genes (DEGs) revealed that ultrasound stimulation induced upregulation of biological processes related to intermediate filament organization, cell–cell signaling, cell adhesion, and epithelial cell differentiation, indicating enhanced structural and developmental maturation (Figure 5C). Gene set enrichment analysis (GSEA) further corroborated these findings. The US group demonstrated significant enrichment of the WNT/β-catenin and mTORC1 signaling pathways compared to the control group, both of which are closely linked to neural development and growth (Figure 5D). In addition, the US group exhibited upregulation of gene sets associated with brain development, accompanied by a consistent downregulation of genes involved in proliferation and cellular damage (Figure 5E-G). Modular network analysis using MCODE identified a core functional cluster enriched for AP-1 transcription factors, notably cFOS and JUNB, together with co-regulators such as SP1 and EGR1. These results highlight the potential of cFOS-centric regulatory circuits as major drivers of gene expression changes induced by ultrasound stimulation in early organoid differentiation (Figure 5H). These findings support the utility of ultrasound as a biophysical modality for the temporal regulation of organoid differentiation and function.

### Ultrasound stimulation enhances the synaptic density of dopaminergic neurons in organoids

In order to comprehensively evaluate the effects of ultrasound stimulation on mBOs, we next investigated synaptic density as a marker of cellular maturation. We immunostained for synaptophysin and postsynaptic density protein 95 (PSD95), which are the presynaptic and postsynaptic markers, respectively (Figure 6A, B). In the case of synaptophysin, there were no significant changes between the control condition and stimulation conditions for MD35 (Figure 6C). On MD55, there was a slight increase in the synaptophysin expression level for the stimulation condition. On the other hand, there was a slight increase in PSD95 levels on MD35 (Figure 6D). On MD55, ultrasound stimulation produced significantly increased levels of PSD95 in the mBOs. These results indicate that ultrasound stimulation enhances synaptic maturation in mBOs, as evidenced by increased PSD95 expression.

**Figure 6.**
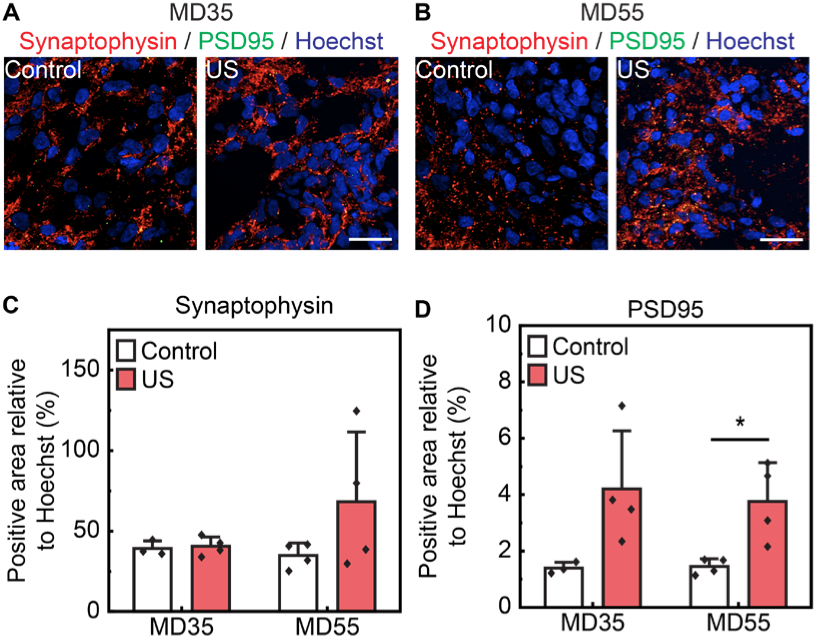
Controlled synaptic density of midbrain organoids using ultrasound stimulation. (A and B) Representative immunostaining images of synaptophysin and PSD95. Scale bar, 50 μm. (C) Relative protein level of synaptophysin in the US group. MD35, two sample two-sided unpaired Student’s t-test, *t=-0.323, p=0.760, n=4*. MD55, two sample two-sided unpaired Student’s t-test, *t=-1.510, p=0.223, n=4*. (D) Relative protein level of PSD95 in the US group. MD35, two sample two-sided unpaired Student’s t-test, *t=–2.694, p=0.072, n=4*. MD55, two sample two-sided unpaired Student’s t-test, *t = −3.283, *p<0.05, n=4*. Solid bars and error bars indicate mean value and standard deviation, respectively.

### Ultrasound stimulation reduces cellular damage markers in organoids

Our stimulation conditions were determined based on established safety guidelines for ultrasound neuromodulation (57, 58). Our mechanical index (MI), and thermal dose were 0.306, and 0.00078 cumulative equivalent minutes (CEM), respectively, which were sufficiently below the FDA and ITRUSST limits (57, 58). The thermal dose was calculated using the measured temperature profile (Figure S12). To experimentally verify the safety of ultrasound stimulation on mBOs, we conducted quantitative analyses of direct and indirect damage markers. First, we measured the diameter of the mBOs on MD2, MD40, and MD55 to examine the effects of ultrasound stimulation on the overall growth. There was no significant change in diameter between the control and stimulated organoids across all measurements, which confirmed that ultrasound stimulation did not inhibit the overall growth of mBOs (Figure S9A, B).

Next, we examined the protein level of apoptotic and DNA damage markers, cleaved caspase 3 (cC3), p53, and phosphorylated H2A.X (Figure 7A-D). For cC3, there was a slight decrease in protein levels on MD35, and a significant decrease on MD55 for the US group (Figure 7E). For p53, there was a slight decrease in protein levels on MD35 for the stimulated group, while on MD55, ultrasound induced a significant decrease in protein levels (Figure 7F). For phosphorylated H2A.X, there was a slight decrease in protein levels for stimulated organoids on both MD35 and MD55 (Figure 7G). These results demonstrated that ultrasound stimulation promoted recovery and neuroprotective effects.

**Figure 7.**
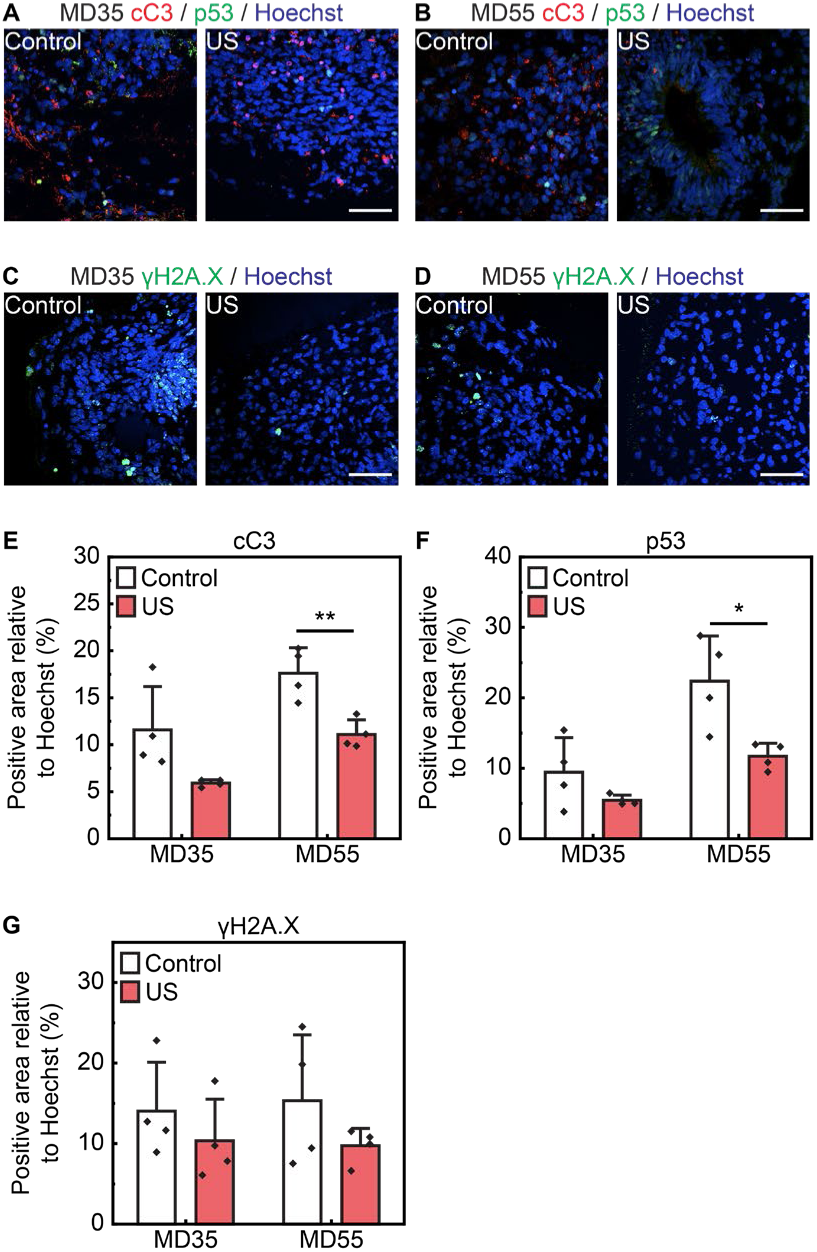
Safety evaluation of ultrasound stimulation on midbrain organoids. (A and B) Representative immunostaining images of cC3 and p53. Scale bar, 50 μm. (C and D) Representative immunostaining images of phosphorylated H2A.X. Scale bar, 50 μm. (E) Relative protein level of cC3 in the US group. MD35, two sample two-sided unpaired Student’s t-test, *t=2.447, p=0.091, n=4*. MD55, two sample two-sided unpaired Student’s t-test, *t=4.179, **p<0.01, n=4*. (F) Relative protein level of p53 in the US group. MD35, two sample two-sided unpaired Student’s t-test, *t=1.604, p=0.204, n=4*. MD55, two sample two-sided unpaired Student’s t-test, *t=3.192, *p<0.05, n=4*. (G) Relative protein level of phosphorylated H2A.X in the US group. MD35, two sample two-sided unpaired Student’s t-test, *t=0.921, p=0.393, n=4*. MD55, two sample two-sided unpaired Student’s t-test, *t=1.327, p=0.266, n=4*. Solid bars and error bars indicate mean value and standard deviation, respectively.

Last, we quantitatively assessed the differentiation of mBO cells into astrocytes by evaluating the expression levels of the S100 calcium-binding protein B (*S100β*) gene and glial fibrillary acidic protein (GFAP). There was no significant difference in both the mRNA expression level of *S100β* and protein expression level of GFAP for the stimulated group on MD35 and MD55, except for a significant increase in the S100β gene expression on MD35 (Figure S13A-D).

## Discussion

Organoids are critical biological models that recapitulate the complexity of organs and organ systems, and precise control of their differentiation timing is essential. Progenitor cells, as precursors to fully differentiated cells, are particularly important for therapeutic applications such as tissue grafting and cell transplantation. In this work, we developed a robust multi-well ultrasound stimulation platform for temporal modulation of midbrain organoid (mBO) differentiation. Our platform was designed for scalability in mind, with design considerations for various well plate sizes. Using a curated protocol and comprehensive biomarker analysis, we demonstrated ultrasound-mediated modulation of the dopaminergic progenitor cell population in mBOs. In addition, we confirmed that ultrasound generally confers a positive effect on mBO growth and maturation.

Using our modular stimulation platform, we first investigated the neuronal response of mBOs to ultrasound stimulation by analyzing cFOS and YAP. As expected, ultrasound successfully activated the cFOS and YAP mechanisms, which is consistent with previously reported studies. Next, we analyzed the dopaminergic progenitor markers FOXA2 and *LMX1A* to assess the effects of ultrasound on the progenitor population. We verified that ultrasound was particularly effective in increasing the dopaminergic progenitor population in the mBOs. Interestingly, *TH* expression and dopamine secretion were reduced at later stages (MD55), suggesting that ultrasound stimulation delays terminal differentiation and temporarily maintains progenitor-like states.

Last, we conducted safety assessments using cC3, p53, and phosphorylated H2A.X markers, which are related to apoptosis and DNA damage. Ultrasound stimulation significantly reduced the expression of all damage markers in the mBOs. Our bulk RNA sequencing results confirmed that the differentiation and maturation of mBOs were largely driven by biochemical cues, with major transcriptomic processes following normal mBO development. The presence of ultrasound stimulation conferred finer control over specific processes related to cellular mechanics. However, we acknowledge that the number of differentially expressed genes (DEGs) was moderate, and the functional interpretation (maturation vs. stress-adaptive pathways) should be viewed as indicative trends that require further validation. Crucially, ultrasound stimulation did not negatively affect mBO growth and temporally modulated the progenitor differentiation stage via a mechanosensitive pathway.

Regarding the increase in the population of dopaminergic progenitor cells and the expression of genes related to brain development in the stimulated group, the activation of YAP is a possible mediator contributing to these changes (59–61). The activation of the JUNB-centered regulatory module, as revealed by network analysis, implicates the involvement of the Hippo and possibly Rho signaling pathways. Given the increased nuclear YAP localization and expression of its downstream target *CTGF*, our findings align with previous reports linking mechanotransduction to neural progenitor regulation. Although further experiments, such as YAP inhibition or pathway-specific perturbations, are required to establish causality, these observations suggest that ultrasound modulates progenitor cell fate via convergence of FOS/JUN, YAP, and cytoskeletal signaling axes.

Nevertheless, our work suggests that ultrasound safely activates neuronal, astrocytic, and differentiation pathways simultaneously without significantly affecting the overall growth, maturation, and function of other cell types within the mBOs. In particular, we demonstrate temporal modulation of progenitor cell differentiation, which is critical for tissue grafting and transplantation. For successful tissue grafting procedures, 2D and 3D stem or progenitor cell culture systems of a specific organ lineage, including 3D progenitor organoids, have been extensively developed and employed in clinical settings (12, 13, 62–64). Such a source of complex differentiated and undifferentiated cells has been traditionally produced using biochemical methods in a free-floating medium. This study presents a potential complementary approach that incorporates ultrasound stimulation to mechanically promote progenitor cell differentiation in an early-stage mBO model.

A limitation of the current study is the lack of investigation into the reversibility or epigenetic nature of the ultrasound-induced effects in the mBOs. However, based on the reported characteristics of LIFU in the literature, we suggest that our stimulation effects are not permanent (32, 65). Further work is necessary to determine the duration of sustained ultrasound-induced effects post-stimulation and whether epigenetic modifications are involved. Nonetheless, our results present a compelling case for using ultrasound stimulation for temporal modulation of differentiation timing in organoids. Future works could build on this study and translate these results to tissue grafting and cell transplantation applications.

## Materials and Methods

### Experimental design

This study aimed to investigate the impact of low-intensity focused ultrasound (LIFU) stimulation on midbrain organoid differentiation and maturation. We hypothesized that LIFU stimulation could modulate the timing of cell-type specific differentiation without inducing cellular damage. A modular ultrasound stimulation platform was designed for multi-well compatibility and delivered ultrasound from maturation day 1 (MD1) to MD55. Post-stimulation, organoids were analyzed using immunostaining, qPCR, ELISA, and bulk RNA sequencing to assess neural activation, dopaminergic progenitor cell differentiation, dopaminergic function, synaptic density, and cellular damage markers.

### Cell culture and generation of midbrain organoids

Human embryonic stem cells (hESCs) from the H9 cell line (WiCell Research Institute, Madison, WI, USA) were cultured in mTeSR-1 medium (STEMCELL Technologies) on Matrigel-coated dishes (Corning). The culture medium was refreshed daily, and cells were passaged using ReLeSR (STEMCELL Technologies) according to the manufacturer’s instructions. hESC experiments were approved by the Korean MoHW Bioethics Committee (IRB No. P01–201409-ES-01) and followed all applicable ethical guidelines.

Midbrain organoids were generated as previously described [PMID: 36121202], with slight modifications. hESCs were dissociated into single cells using Accutase and seeded at a density of 8 × 10³ cells per well in ultra-low attachment 96-well plates to initiate self-organization into embryoid bodies. Upon formation of embryoid bodies, the culture medium was changed to N2 medium composed of a 1:1 mixture of DMEM/F12 and Neurobasal medium supplemented with 1× N2 supplement (Gibco), 1× β-mercaptoethanol, 1% non-essential amino acids (NEAA), 1% penicillin-streptomycin (Pen/Strep), and 1% GlutaMAX. This medium was further supplemented with 10 nM SB431542, 150 ng/ml Noggin, 1.5 μM CHIR99021, and 400 ng/ml sonic hedgehog (SHH), and replaced on days 2, 4, and 6.

On day 8, the medium was changed to N2 medium containing 100 ng/ml fibroblast growth factor 8 (FGF8). On day 11, the culture medium was switched to B27 medium consisting of Neurobasal medium supplemented with 1× B27 supplement without vitamin A (Gibco), 1× β-mercaptoethanol, 1% NEAA, 1% Pen/Strep, and 1% GlutaMAX, further enriched with 100 ng/ml FGF8b, 20 ng/ml brain-derived neurotrophic factor (BDNF), and 200 μM ascorbic acid. From day 14 onward, the organoids were maintained in maturation medium composed of B27 medium supplemented with 20 ng/ml BDNF, 10 ng/ml glial cell-derived neurotrophic factor (GDNF), 200 μM ascorbic acid, 500 μM cyclic AMP (cAMP), and 1 μM DAPT, with fresh factors replenished regularly.

### Ultrasound stimulation platform

The modular PCB was designed (Autodesk Fusion 360, Autodesk, Inc., USA) and fabricated (R&D Tech Co., Ltd, Republic of Korea) for a single-element transducer. A circular lead zirconate titanate (PZT) disk (H4P161000, PZT Electronic Ceramic Co., Ltd, China) with a diameter of 16 mm and a thickness of 2 mm was soldered onto the PCB using a low-temperature solder (SMD291AX, Chip Quik Inc., Canada). An SMA connector (132165, Amphenol RF, USA) was also soldered on the PCB for interfacing with the ultrasound driving circuitry. Then, the assembled PCB was dip-coated with PDMS, which was prepared by mixing the base and curing agent at a 10:1 ratio, and the PZT disk was covered with a polymethylpentene film (PMP, TPX® DX845, Goodfellow Cambridge Ltd., England) for increased robustness. The PDMS was cured on a hotplate at 90°C for at least 4 hours. To evaluate the robustness of the modular platform, the electrical impedance was measured (E4990A, Agilent Technologies Inc., USA) before stimulation and after every other stimulation period for a total of 9 full operation cycles. All measurements were conducted after 6 hours of operation in the incubator.

### FEM simulation of ultrasound beam

We simulated the ultrasound beam profile for platforms with different PZT diameters, culture well size, and vibration modes of PZT using FEM (COMSOL Multiphysics, MA, USA). To minimize the computation time, we simulated in axisymmetric two dimensions (2D). The geometry of each component, such as culture well, and medium, was set to the same actual size and location. We selected PZT disks with diameters closely matching the bottom size of each well plate. The diameters were set to 35 mm, 16 mm, and 8 mm for 6-well, 24-well, and 96-well plates, respectively.

We applied Solid Mechanics, Electrostatics, and Pressure Acoustics, Frequency Domain physics modules. Moreover, the Piezoelectric Effect physics derived from Solid Mechanics and Electrostatics, and Acoustic-Structure Boundary derived from Pressure Acoustics, Frequency Domain, and Solid Mechanics were also added. We applied the terminal voltage at the bottom surface of the PZT disk and grounded the top surface of the PZT disk. We solved these systems with a frequency domain study using a frequency of radial mode operation in 96-well, 24-well, and 6-well systems, respectively, and thickness mode operation in 24-well systems. We exported the acoustic intensity magnitude profile into CSV format and processed it with MATLAB software. Using MATLAB software, we plotted a simulated beam profile and 50% intensity contours. We also calculated FWHM, the ratio of the 50% intensity volume within the organoid space, and maximum intensity variation within the organoid space.

### Ultrasound beam profile measurement

We characterized the ultrasound beam profile of the fabricated platform using a hydrophone (NH0500, Precision Acoustics, UK). The XYZ-location of the hydrophone is precisely controlled by a customized motor system (Sciencetown Inc., Republic of Korea) and a custom-made MATLAB program (Mathworks®, USA) (41, 50). The transducer platform was attached beneath the culture well with ultrasound transmission gel. Approximately 500 μL of soybean oil was filled in the culture well, and the hydrophone measured the ultrasound beam profile in the oil by sweeping a cubic volume of 8 mm × 8 mm × 2 mm. The measured voltage signal was amplified and DC-coupled (Precision Acoustics, UK), and the modified signal was acquired with a digital oscilloscope (DSOX2022A, Agilent Technologies, USA). The obtained voltage values were converted to I_SPPA_ values by using the following formula.

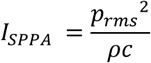

where *p_*r*ms_* is the root-mean-square pressure, *ρ* is the density of the media, and *ρ* is the speed of sound in the media.

### Ultrasound stimulation equipment and protocol

We stimulated midbrain organoids under ultrasound conditions and a control condition. The fabricated ultrasound stimulation platform was operated using a function generator (33220A, Agilent Technologies, USA). The ultrasound stimulation was conducted from MD1 to MD55. No ultrasound stimulation was delivered for the control condition.

### Thermal characteristics of the stimulation platform

To investigate the heat generation due to ultrasound, we examined the temperature change at the bottom of the well plate. We used the same setup as stimulation, the same well plate, and the same amount of media. The well plate with media was put inside the incubator, and the tip of the thermocouple (T9234, Daihan Scientific Co., Korea) was located at the center of the well plate bottom. Before operating the PZT, we waited for the temperature to stabilize over a period of 10 minutes. We operated the PZT using the US group condition used for the main *in vitro* experiments, and the temperature was measured at 1-minute intervals over 20 minutes. MI and CEM are determined by the following formulas:

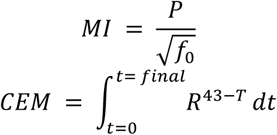

where *P* is the pressure amplitude, *f*_0_ is the operating frequency, *d* is the time for the temperature change, and *T* is the temperature. *T* was set using the measured profile (Figure S10), and *R* was set at 0.25 for *T* < 43℃ (57, 58).

### Immunostaining

For MD1 mBOs, organoids were fixed in 4% paraformaldehyde (PFA) at room temperature for 6 hours and subsequently washed with 1× PBS. Fixed organoids were embedded in optimal cutting temperature (OCT) compound (Tissue-Tek®; Sakura Finetek, USA) and cryosectioned into 7 μm-thick slices using a cryostat (CM1520, Leica, Germany). For immunostaining, sections were permeabilized in 0.3% Triton X-100 (Sigma) in 0.025% TBS-T for 1 hour, followed by blocking with 3% bovine serum albumin (BSA) and 0.02% sodium azide in 0.025% TBS-T for 1 hour at room temperature. Sections were incubated overnight at 4 °C with primary antibodies diluted in blocking buffer: anti-SOX2 (Seven Hills Bioreagents, #WRAB1236), anti-tyrosine hydroxylase (TH; Sigma, #T1299), and anti-MAP2 (R&D Systems, #MAB933). After washing with 1× TBS-T, sections were incubated for 1 hour at room temperature in the dark with Alexa Fluor-conjugated secondary antibodies (Alexa Fluor 488 goat anti-mouse IgG and Alexa Fluor 555 donkey anti-rabbit; Life Technologies) along with Hoechst nuclear stain (Thermo Scientific™). Fluorescent images were acquired using a confocal laser scanning microscope (LSM800, Zeiss, Germany). Excluding MD1 mBOs, organoids were fixed in 4% paraformaldehyde at room temperature for 5 hours. Stimulated organoids were fixed 1 hour after the end of stimulation. Then, organoids were washed 3 times with phosphate-buffered saline (PBS), each for 30 minutes. Washed organoids were immersed overnight in a 30% sucrose solution for cryoprotection. Then, the organoids were embedded in an optimal cutting temperature (OCT) compound (Tissue-Tek®; Sakura Finetek, USA) to produce the frozen blocks. Frozen organoids were sectioned into 7-μm-thick slices using a cryocut microtome (CM3050s, Leica, Germany), and sections were obtained from three distinct depths. After drying overnight, sliced samples were washed 3 times with PBS, each for 5 minutes, and permeabilized with 0.3% Triton X-100 in Tris-buffered saline containing 0.025% Tween 20 (TBS-T) solution for 50 minutes. The nonspecific binding of antibodies was blocked with a blocking solution, which contained 3% bovine serum albumin (BSA) and 0.02% sodium azide in 0.025% TBS-T solution for 30 minutes. Primary antibodies were diluted with the blocking solution: anti-phospho-Histone H2A.X (Sigma Aldrich, #05-636), anti-MAP2 (R&D Systems, #MAB8304), anti-p53 (Santa Cruze Biotechnology, #sc-126), anti-PSD95 (Invitrogen, #MA1-046), anti-cC3 (Cell Signaling Technology, #9664s), anti-cFOS (Abcam, ab190289), anti-FOXA2 (Seven Hills Bioreagents, #WRAB1200), anti-GFAP (DAKO, #Z0334), anti-synaptophysin (Abcam, #ab32127), anti-TH (Novus Biologicals, #NB300-109), anti-YAP (Cell Signaling Technology, #14074). Samples were incubated overnight with diluted primary antibodies at 4℃. Next, samples were washed 3 times with PBS, each for 5 minutes, and incubated with secondary antibodies (Cy3 goat anti-Rabbit IgG; Jackson, Alexa Fluor 555 donkey anti-rabbit; Life Technologies, Alexa Fluor 488 goat anti-mouse IgG; Invitrogen, Alexa Fluor 488 goat anti-mouse IgG; Life Technologies) and Hoechst (Thermo Scientific™) for 1 hour at room temperature in the dark. Again, samples were washed 3 times with PBS for 5 minutes each. The slide glass with stained samples was covered with a coverslip after dropping the mounting solution. Stained samples were observed under a confocal microscope (LSM980, Zeiss, Germany / AX, Nikon, Japan) and analyzed with ImageJ software (National Institutes of Health, Bethesda, MD, United States).

### Real-time PCR

Organoids were washed with dulbecco’s phosphate-buffered saline (dPBS) for 30 min, 1 hour after the end of stimulation. Total RNA was extracted from organoids using the easy-BLUETM Total RNA Extraction Kit (iNtRON Biotechnology) following washing with 1× PBS. RNA was purified through chloroform and isopropanol precipitation, and the resulting RNA pellet was washed with cold 70% ethanol. RNA concentration was determined using a NanoDrop spectrophotometer (Thermo Fisher Scientific). A total of 1200 ng RNA was reverse transcribed into complementary DNA (cDNA) using the SuperScript IV Reverse Transcriptase kit (Thermo Fisher Scientific). Quantitative real-time PCR was performed using Fast SYBRTM Green PCR Master Mix (Applied Biosystems) and gene-specific primers to evaluate marker gene expression levels.

### Dopamine ELISA

Cultured medium was collected 24 hours after incubation from each well of a 24-well plate, containing a single organoid. Collected medium was analyzed using a dopamine ELISA kit (ENZ-KIT188, Enzo, Switzerland) according to the instructions of the manufacturer. Each sample was added to a plate well, and the same volume of biotin-detection antibody working solution was added. The plate was incubated for 45 minutes at 37℃ and washed. After washing, horseradish peroxidase (HRP)-streptavidin working solution was added into each well and incubated for 30 minutes at 37℃. The plate was washed again, and tetramethylbenzidine (TMB) substrate was added. The plate was then incubated in the dark for 15 minutes at 37℃. After incubation, a stop solution was added immediately. The optical density absorbance at 450 nm was measured by a microplate reader (#EPOCH-SN, Agilent, CA, United States). Dopamine concentration of samples was calculated with a standard curve using GraphPad Prism software 6.0 (GraphPad Software, Inc., CA, United States).

### Bulk RNA sequencing

To comprehensively assess gene expression profiles in midbrain organoids, bulk RNA sequencing was performed by Macrogen (Seoul, Korea). Samples were collected from four experimental groups, including PSC, mBO-control, US at day 35 of differentiation (MD35). RNA libraries were prepared using the TruSeq Stranded mRNA LT Sample Prep Kit (Illumina, San Diego, CA, USA), and sequencing was conducted on an Illumina platform. Raw sequencing reads were subjected to quality trimming and aligned to the human reference genome (GRCh38) using HISAT2. Transcript assembly was carried out with StringTie, and transcript abundance was quantified in fragments per kilobase of transcript per million mapped reads (FPKM). Expression data were further normalized by log₂ transformation [log₂(FPKM + 1)] prior to downstream analyses. Gene set enrichment analysis (GSEA) was conducted using GSEA Desktop v4.3.3 (Broad Institute) with all type signature gene sets from the Human MSigDB Collections (v2024). Pearson correlation coefficients (r) between samples from each group (control, US) were independently calculated using ExDEGA v5.2.1. Differentially expressed genes (DEGs) were filtered based on a fold-change threshold of 2 on PSC-mBO comparison and >1.5 or <0.7 on ultrasound stimulation comparison and visualized by heatmap generation in MeV 4.9.0. Pathway enrichment analyses were conducted using the DAVID Bioinformatics Resources v6.8 (https://david.ncifcrf.gov) and the Reactome database (https://reactome.org/), including MCODE. Resulting pathway networks were visualized and further analyzed using Cytoscape software (www.cytoscape.org).

### Statistical analysis

Normal distribution of all *in vitro* data sets was confirmed using the Shapiro-Wilk test. For immunohistology data and RT-PCR data, a two-sample two-sided unpaired Student’s t-test was conducted on normally distributed data. For non-parametric data, the Mann-Whitney U test was used. For ELISA data, a one-sample two-sided Student’s t-test against a value of 1 was conducted. All data were analyzed using Microsoft Excel (Microsoft Co., WA, USA), OriginPro 2019 (OriginLab Co., MA, USA), and MATLAB R2022b (Mathworks, MA, USA). Unless stated otherwise, all data are presented as mean ± standard deviation (SD), and a p-value of <0.05 was considered significant. Graphical illustrations of biological components in all relevant figures were created with Adobe Illustrator 2023 and BioRender.com.

## Acknowledgments

We would like to acknowledge the facilities and the scientific and technical assistance of the KARA (KAIST Analysis Center for Research Advancement), KAIST Bio Core Center, and Microscopy and Imaging Analysis Core Facility in the BioMedical Research Center (BMRC), KAIST. We also thank S. Bang for his comments on the graphical illustrations.

## Funding

K-Brain Project of the National Research Foundation (NRF) funded by the Korean government (MSIT) RS-2023-00262568 (HJL)

Korea Dementia Research Project through the Korea Dementia Research Center (KDRC), funded by the Ministry of Health & Welfare and Ministry of Science and ICT, Republic of Korea RS-2024-00355871 (HJL)

BK21 FOUR Connected AI Education & Research Program for Industry and Society Innovation, KAIST EE, No. 4120200113769 (HJL)

Korea-US Collaborative Research Fund(KUCRF), funded by the Ministry of Science and ICT and Ministry of Health & Welfare, Republic of Korea RS-2025-16022980 (HJL)

Korea Research Institute of Bioscience and Biotechnology (KRIBB) Research Initiative Program KGM4722533 (M-OL)

## Supporting Information

**Figure S1.**
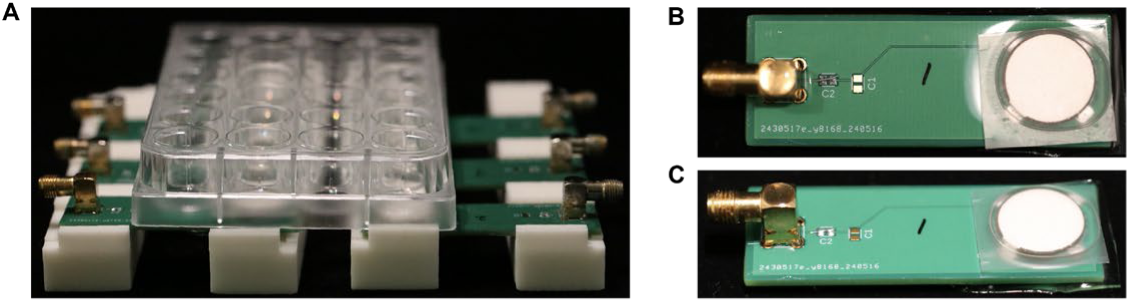
Photos of ultrasound transducer platforms. (A) Integration of 24-well plates and six ultrasound transducer platforms placed inside the incubator. (B) Top view of a single ultrasound transducer platform. (C) Front view of the ultrasound transducer platform.

**Figure S2.**
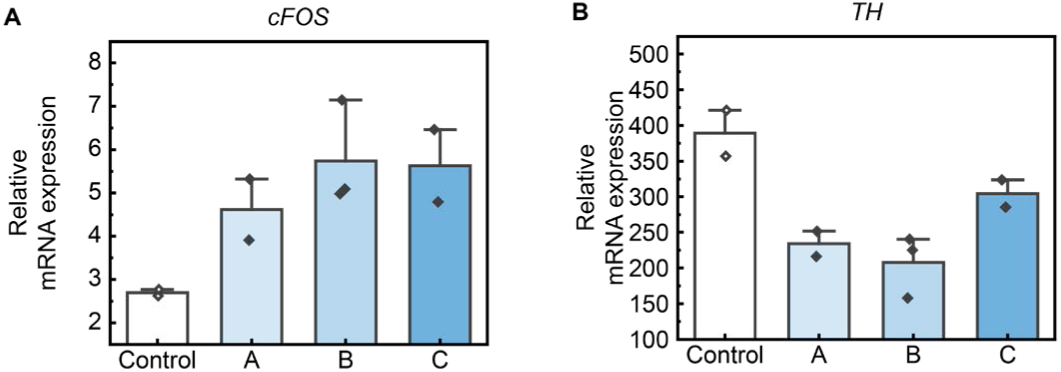
Preliminary data for PRF selection. (A) Relative mRNA expression of *cFOS* across three different PRFs. (B) Relative mRNA expression of *TH* across three different PRFs. Solid bars and error bars indicate mean value and standard deviation, respectively.

**Figure S3.**
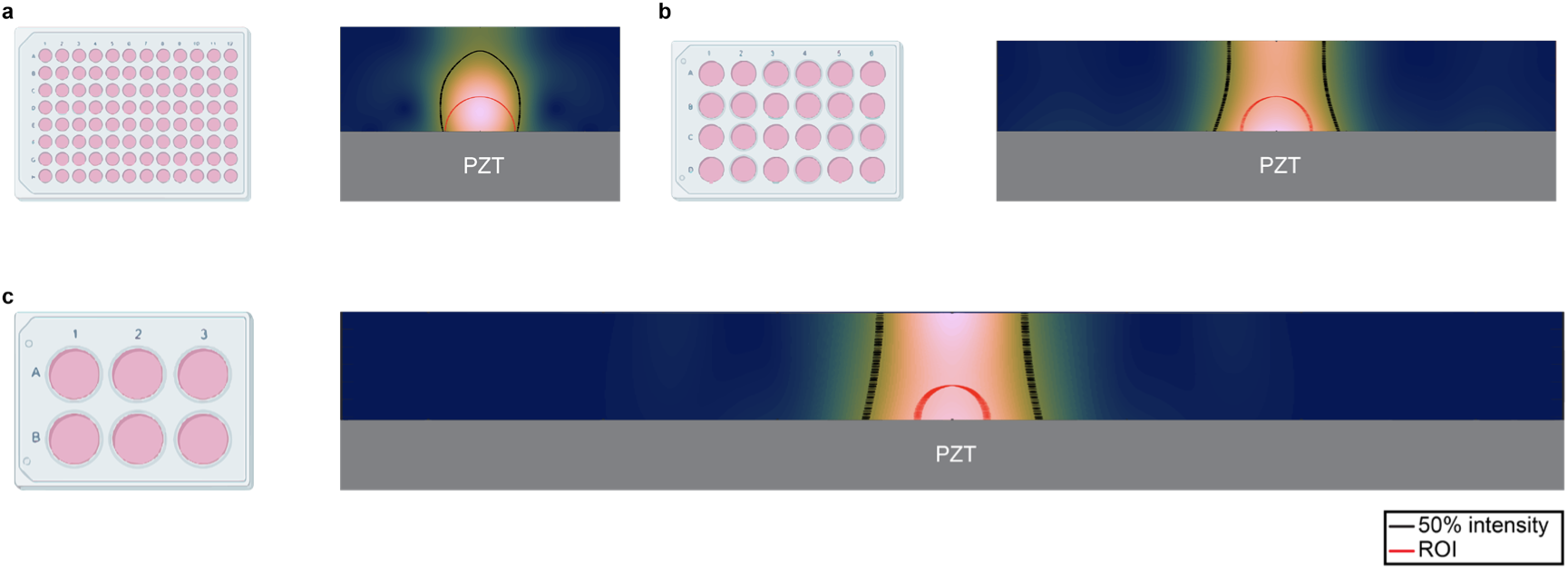
FEM simulation of the ultrasound beam for optimizing the Well-PZT size. (A) Single well simulation results of a 96-well plate. (B) Single well simulation results of a 24-well plate. (C) Single well simulation results of a 6-well plate.

**Figure S4.**
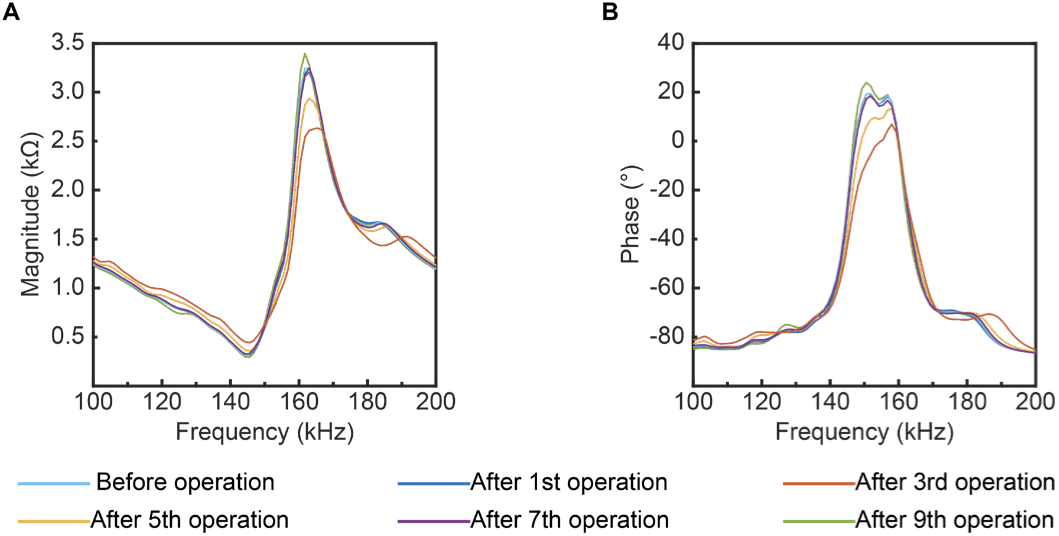
Robustness test of the platform over 9 operations. (A) Magnitude of measured electrical impedances. (B) Phase of measured electrical impedances.

**Figure S5.**
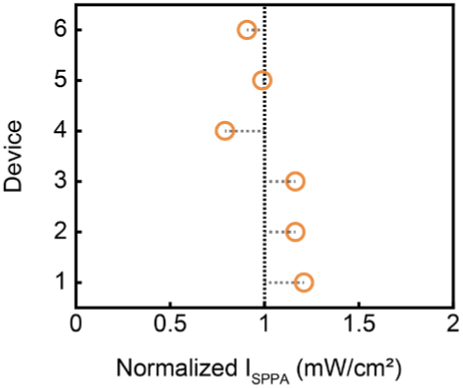
Evaluation of ultrasound intensity variations across six platforms. Normalized I_SPPA_ of each platform for generating ultrasound.

**Figure S6.**
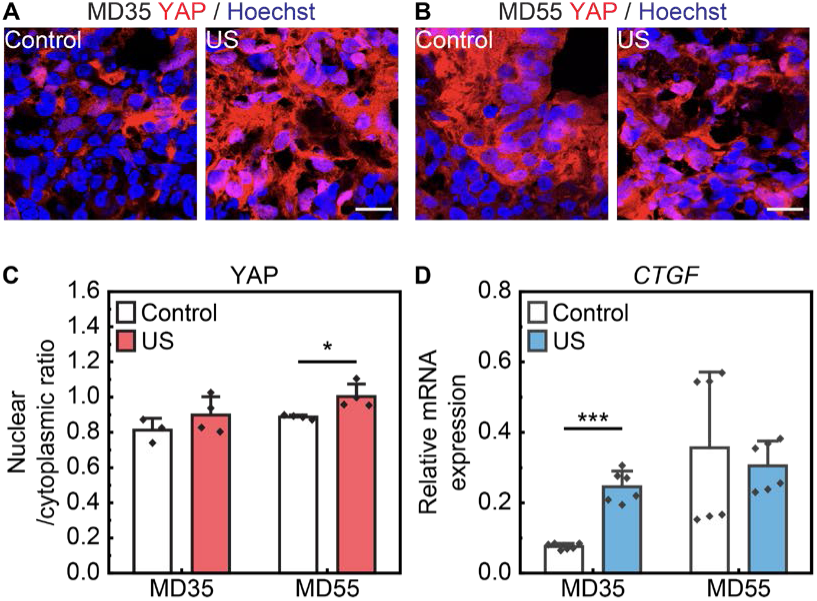
Additional analysis of ultrasound-induced cellular activation. (A and B) Representative immunostaining images of YAP. Scale bar, 20 μm. (C) Nuclear translocation ratio of YAP in the US group. MD35, two sample two-sided unpaired Student’s t-test, *t=–1.338, p=0.239, n=4*. MD55, two sample two-sided unpaired Student’s t-test, *t=–3.245, *p<0.05, n=4*. (D) Relative mRNA expression of *CTGF* in the US group. MD35, two sample two-sided unpaired Student’s t-test, *t=–9.235, ***p<0.001, n=6*. MD55, Mann-Whitney U test, *U=18, p=1, n=6*. Solid bars and error bars indicate mean value and standard deviation, respectively.

**Figure S7.**
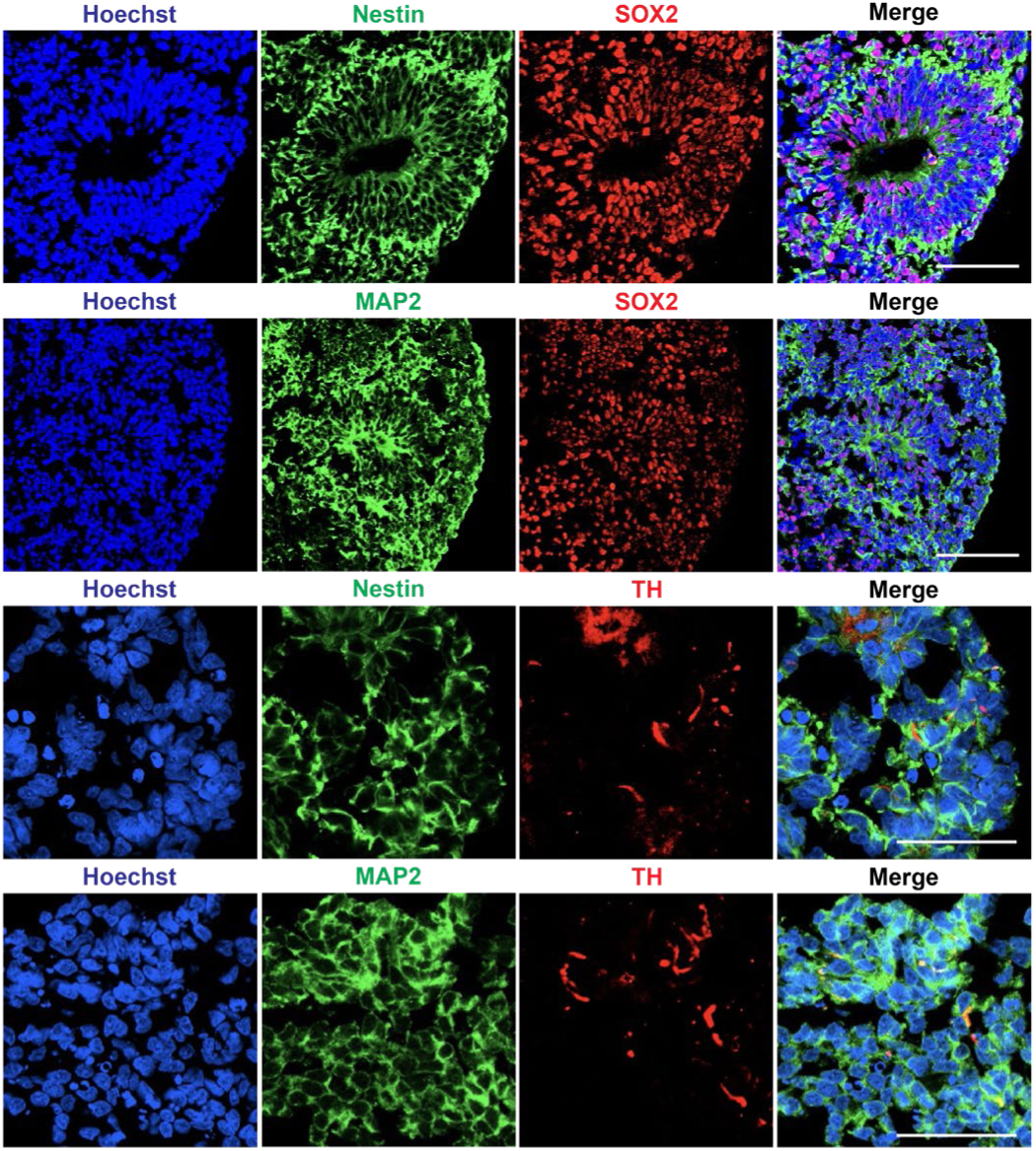
Characterization of MD1 midbrain organoids. Representative immunofluorescence images demonstrating the expression of neural stem cell marker (Nestin), stemness marker (SOX2), neuron marker (MAP2), and dopaminergic neuron marker (TH). Scale bar, 50 μm

**Figure S8.**
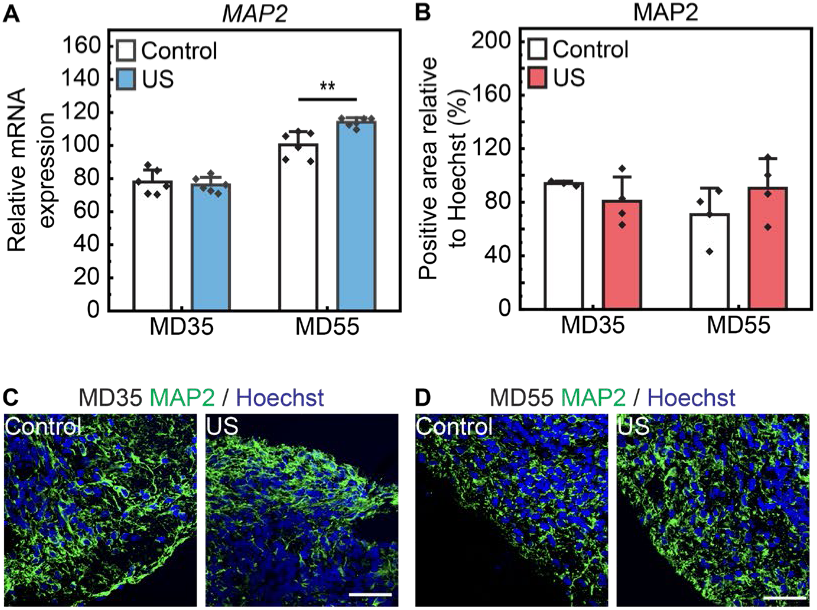
Effects of ultrasound stimulation on neuronal differentiation. (A) Relative mRNA expression of *MAP2* in the US group. MD35, two sample two-sided unpaired Student’s t-test, *t=0.519, p=0.617, n=6*. MD55, two sample two-sided unpaired Student’s t-test, *t=-3.988, **p<0.01, n=6*. (B) Relative protein level of MAP2 in the US group. MD35, two sample two-sided unpaired Student’s t-test, *t=1.452, p=0.240, n=4*. MD55, two sample two-sided unpaired Student’s t-test, *t=-1.320, p=0.236, n=4*. (C and D) Representative immunostaining images of MAP2. Scale bar, 50 μm. Solid bars and error bars indicate mean value and standard deviation, respectively.

**Figure S9.**
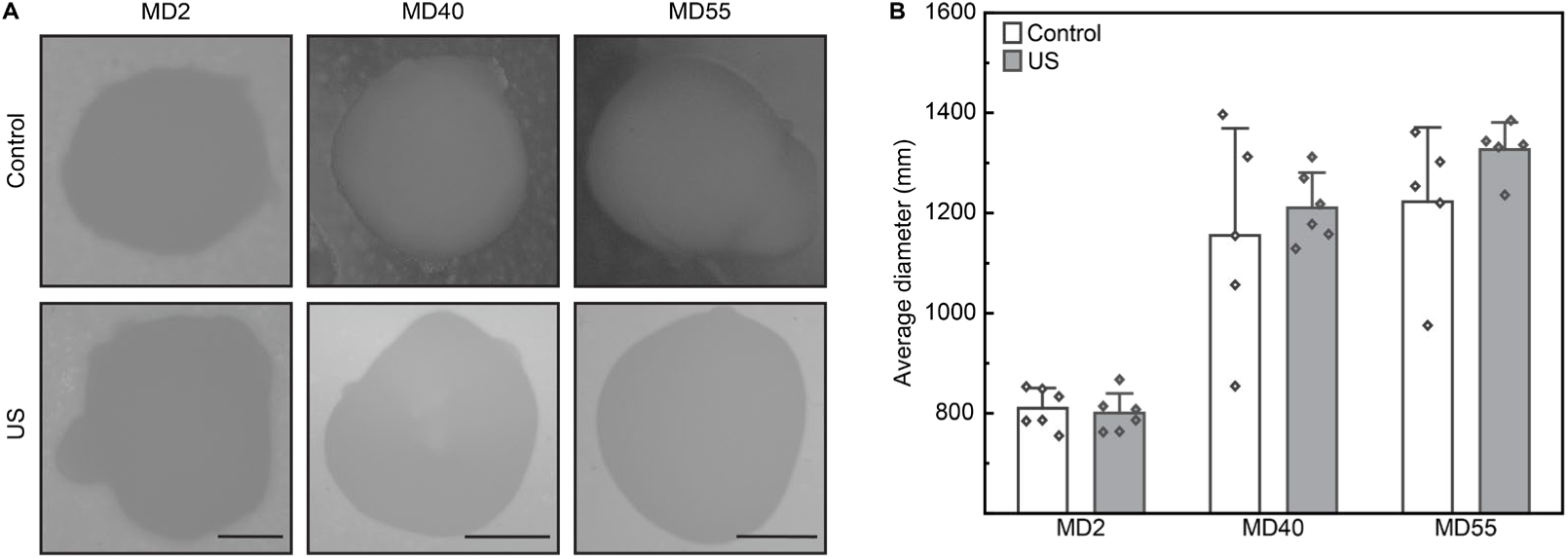
Effects of ultrasound stimulation on the growth of midbrain organoids. (A) Widefield images of midbrain organoids. Scale bar, 250 μm (MD2), 500 μm (MD40), and 500 μm (MD55). (B) Size of the midbrain organoids. Solid bars and error bars indicate mean value and standard deviation, respectively.

**Figure S10.**
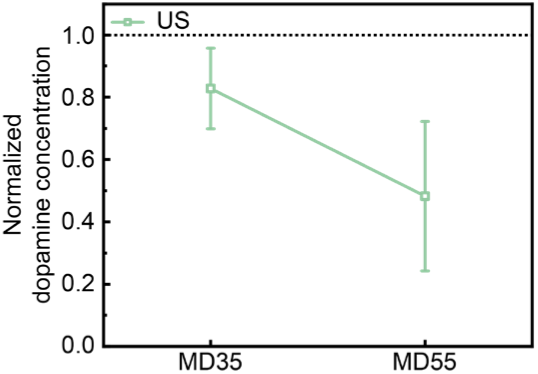
Analysis of organoid dopamine secretion. Dopamine concentrations were normalized to the control group. On MD35, one sample Student’s t-test, *t=-2.295, p=0.149, n=3*. On MD55, one sample Student’s t-test, *t=-3.739, p=0.065, n=3*. Error bars indicate standard deviation.

**Figure S11.**
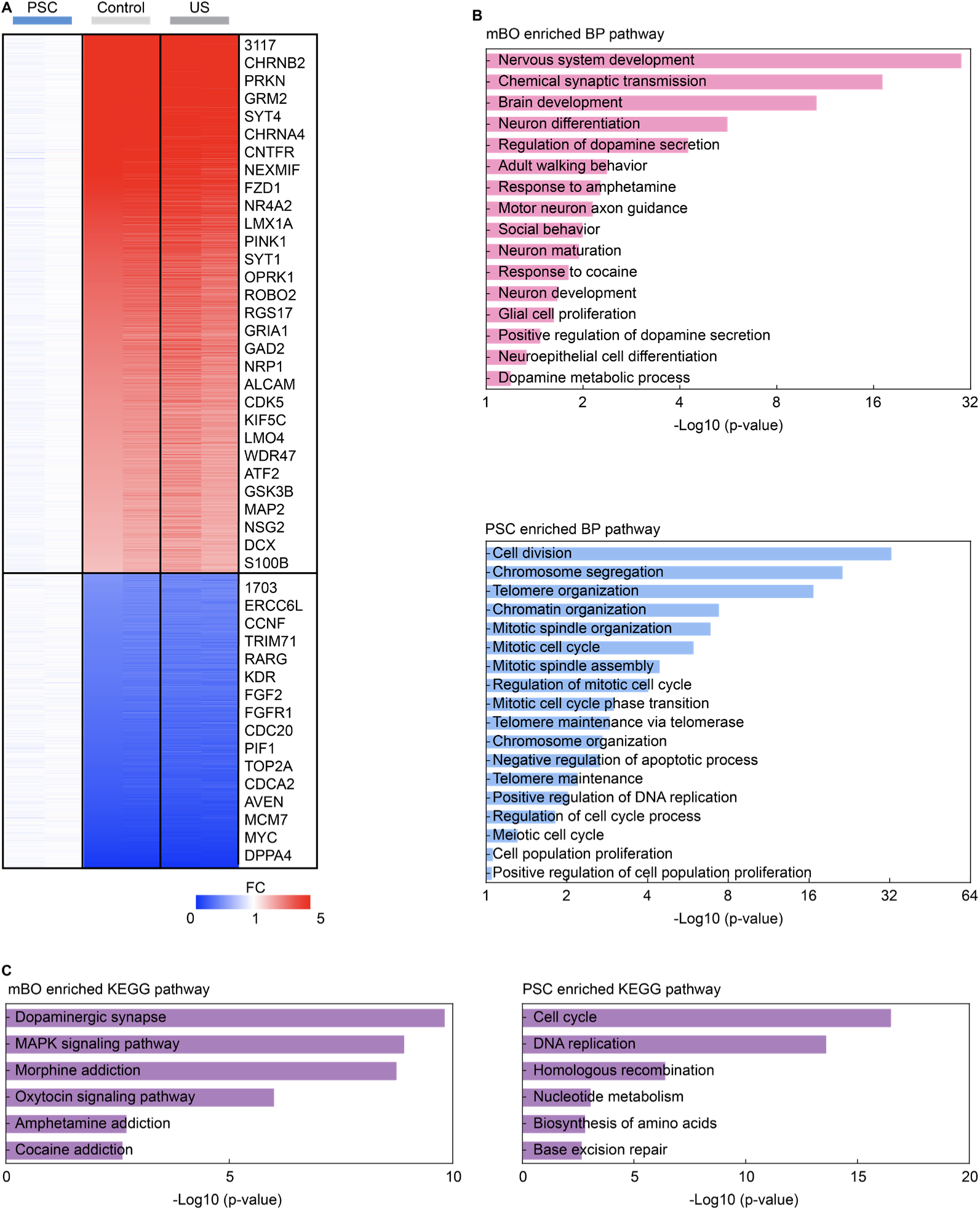
Total transcriptome analysis of PSCs. (A) Heatmap plot for DEGs. (B) GO analysis of the PSCs and mBOs. (C) KEGG pathway analysis of the mBOs and PSCs..

**Figure S12.**
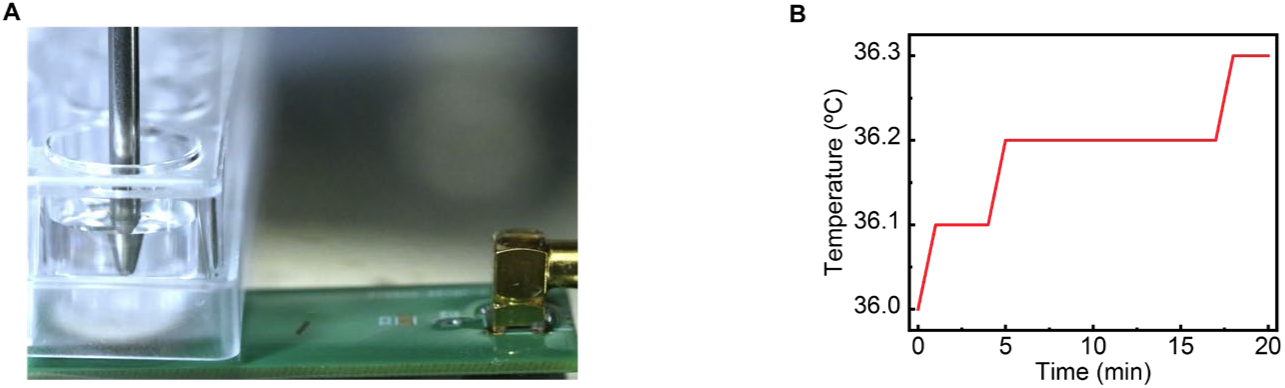
Measurement of transient temperature changes at the surface of the ultrasound transducer platform. (A) Temperature measurement setup in the incubator. (B) Temperature measurement results over 20 minutes.

**Figure S13.**
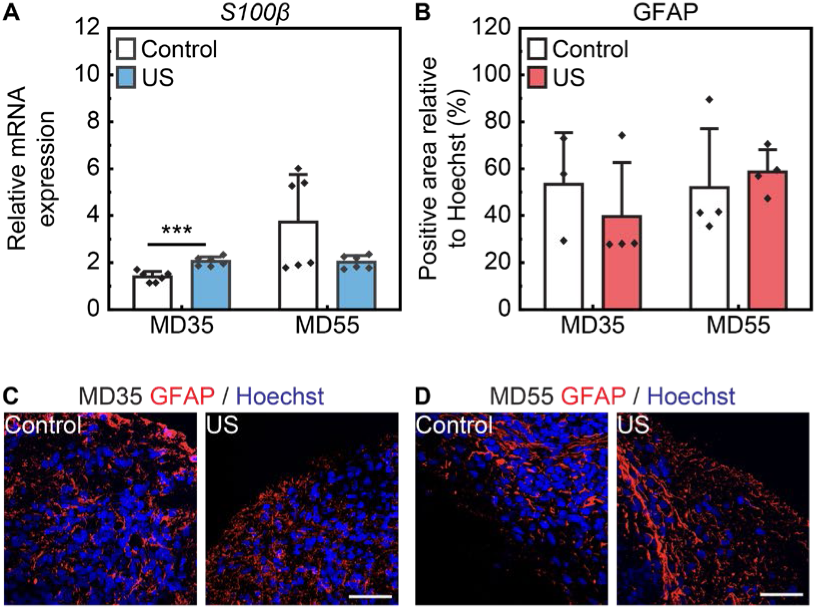
Effects of ultrasound stimulation on astrocytic differentiation. (A) Relative mRNA expression of *S100β* in the US group. MD35, two sample two-sided unpaired Student’s t-test, *t=-5.420, ***p<0.001, n=6*. MD55, Mann-Whitney U test, *U=26, p=0.240, n=6*. (B) Relative protein level of GFAP in the US group. MD35, Mann-Whitney U test, *U=9, p=0.4, n=4*. MD55, Mann-Whitney U test, *U=4, p=0.343, n=4*. (C and D) Representative immunostaining images of GFAP. Scale bar, 50 μm. Solid bars and error bars indicate mean value and standard deviation, respectively.

**Dataset S1 (separate file).** Raw data file for ICC analysis of MD35 organoids

**Dataset S2 (separate file).** Raw data file for ICC analysis of MD55 organoids

